# Standardizing the determination and interpretation of *P*_crit_ in fishes

**DOI:** 10.1101/703991

**Authors:** Jessica E. Reemeyer, Bernard B. Rees

## Abstract

For most fishes, there is an oxygen level, the critical oxygen tension (*P*_crit_), below which oxygen consumption (*M*_O2_) becomes dependent upon ambient oxygen partial pressure (*P*_O2_). We compare multiple curve-fitting approaches to estimate *P*_crit_ of the Gulf killifish, *Fundulus grandis*, during closed and intermittent-flow respirometry. The traditional approach fitting two line segments of *M*_O2_ versus *P*_O2_ produced high and variable estimates of *P*_crit_. Nonlinear regression using hyperbolic or Weibull functions resulted in either variable *P*_crit_ estimates or, in some cases, failed to converge upon meaningful solutions. *P*_crit_ determined as the *P*_O2_ when *M*_O2_ equals standard metabolic rate (SMR) based upon a linear relationship of *M*_O2_ and *P*_O2_ at low *P*_O2_ were consistent across fish and experimental trials. Therefore, we recommend that Pcrit specifically refer to the *P*_O2_ below which SMR cannot be maintained. Its determination, therefore, requires accurate measurement of SMR.

## INTRODUCTION

There is considerable interest in describing the oxygen dependence of aerobic metabolism of animals, especially for animals from aquatic habitats where the oxygen concentration is much lower and more variable than in terrestrial habitats. Determination of this oxygen dependence is particularly relevant in the current context of human-induced environmental change, where increased nutrient input, warmer temperatures, and changes in hydrology have increased the geographic scope and severity of aquatic hypoxia (Diaz and Rosenberg 2008, Rabalais et al., 2010).

Perhaps the most common metric of the oxygen dependence of aerobic metabolism is the critical oxygen tension, *P*_crit_. For an animal that is capable of regulating its metabolism over a broad range of oxygen levels (an oxy-regulator), *P*_crit_ represents the *P*_O2_ where oxygen consumption (*M*_O2_) switches from being independent to being dependent on *P*_O2_ with further decreases in ambient oxygen (Ultsch et al., 1981; Rogers et al., 2016; Wood, 2018). Alternatively, *P*_crit_ has been defined as the *P*_O2_ below which an animal’s basic metabolic needs, i.e. standard metabolic rate (SMR) in fishes, can no longer be sustained aerobically (Claireaux and Chabot, 2016; Thuy et al., 2010; Pan et al., 2016; Snyder et al., 2016; Wong et al., 2017). This level of oxygen was originally described by Fry and Hart (1948) as the “level of no excess activity”. Although related, these two concepts of *P*_crit_ differ because the former depends upon the intensity of metabolism, whereas the latter applies to the level of oxygen that limits a specific metabolic state (Claireaux and Chabot, 2016).

Recently, Wood (2018) questioned the usefulness of the *P*_crit_ concept based on two main concerns: uncertainty of its biological meaning and lack of standardization in its determination. The purpose of this study is not to argue the biological relevance of *P*_crit_, as this concern has been addressed (Regan et al., 2019): rather, the purpose of this study is to evaluate analytical methods used to determine *P*_crit_ from respirometric data. Traditionally, *P*_crit_ has been estimated the intersection of two straight lines, one fit to a region where *M*_O2_ is relatively independent of *P*_O2_ and a second line describing the decrease in *M*_O2_ at low *P*_O2_ (Yeager and Ultsch 1989; Rogers et al., 2016). Because respirometric data rarely conform neatly to two straight lines across a broad range of *P*_O2_, alternative linear or nonlinear regression solutions to determine *P*_crit_ have been proposed (Marshall et al., 2013; Claireaux and Chabot, 2016; Cobbs and Alexander, 2018). Here, we measured *M*_O2_ as a function of *P*_O2_ in closed and intermittent respirometry with the Gulf killifish, *Fundulus grandis*, and applied multiple curve-fitting methods to estimate *P*_crit_. Based upon our results, we recommend that *P*_crit_ be determined as the *P*_O2_ at which *M*_O2_ equals SMR, which can be done with simple linear regression of *M*_O2_ versus *P*_O2_ at low *P*_O2_ (Claireaux and Chabot, 2016). For this method to be general and reproducible, it is imperative that SMR be accurately determined (Chabot et al., 2016).

## MATERIALS AND METHODS

### Animals

Adult male *F. grandis* (n=11) were purchased from local bait shops in the summer of 2018 and housed at the University New Orleans under a 12:12 (light:dark) photoperiod in aerated, filtered 1/3 strength seawater (salinity ≈ 10) at ~27 °C. Fish were fed an amount of flake fish food equal to 1 – 1.5% of their body mass once per day. Fish were identified by unique PIT tags (Reemeyer et al., 2019) or housed individually. Fish were maintained under these conditions for at least one month before experiments. All procedures were approved by the University of New Orleans Institutional Animal Care and Use Committee (Protocol # 18-006).

### Respirometry

Each fish was used in a sequence of three respirometry trials. Trials 1 and 2 employed intermittent respirometry to estimate SMR and RMR (Svendsen et al., 2016), followed by closed respirometry to estimate *P*_crit_. In Trial 3, neither SMR nor RMR was determined, and *P*_crit_ was determined by intermittent respirometry. Trials were separated by approximately one week and they were performed at 27.0 ± 0.5°C in 1/3 strength seawater. Oxygen consumption (*M*_O2_) by fish was determined as previously described (Reemeyer et al., 2019) and outlined below. Fish were starved for 24 h prior to respirometry.

For Trials 1 and 2, fish were weighed (to the nearest 0.01 g) and placed into respirometry chambers between 14:00-15:00. For the first hour, the following intermittent respirometry protocol was used: 60 s flush; 30 s wait; and 120 s *M*_O2_ measurement. At that point, the protocol was adjusted to 300 s flush, 60 s wait, and 240 s *M*_O2_ measurement, which was continued for approximately 14 h. Throughout the combined ~15 h period, *P*_O2_ was maintained at >85% of the air-saturated value. At 06:00 the following morning, the flush pumps were turned off. At that point, the chambers, recirculating pumps, and oxygen sensors formed closed systems, and the *P*_O2_ declined due to *M*_O2_ by the fish. During the closed period, *M*_O2_ was measured over consecutive 60 s intervals until the fish were unable to maintain equilibrium for ≥ 3 s. At that point, the flush pumps were turned on to reoxygenate the chambers. The total time the chambers remained closed ranged from 45 and 108 min. All fish recovered upon reoxygenation, whereupon they were returned to their holding tank.

For Trial 3, fish were weighed (to the nearest 0.01 g) and placed in respirometry chambers between 15:00 and 16:00. Chambers were flushed continuously with well aerated water (> 95% air-saturation) until 21:00. At that time, the *P*_O2_ was stepped down at 1 h intervals by introducing nitrogen gas via a computer-controlled solenoid valve. Target values of *P*_O2_ were 20.75 kPa, 13.07 kPa, 8.30 kPa, 5.19 kPa, 3.32 kPa, 2.07 kPa. Over the last 30 min at each *P*_O2_, *M*_O2_ was measured in three cycles of 300 s flush, 60 s wait, 240 s measurement. Trials ended around 03:00, after which the water was reoxygenated with air. After 30 min recovery, fish were returned to their holding tanks. Importantly, all *P*_crit_ determinations were done during the dark phase of the photoperiod. The only illumination was that required to operate the computer (e.g., to start a closed respirometry trial or to activate nitrogen gassing in the intermittent trials), from which fish chambers were shielded.

*M*_O2_ due to microbial respiration was measured before and after each trial and the *M*_O2_ by each fish was corrected by subtracting a time-weighted background respiration (Reemeyer et al., 2019; Rosewarne et al., 2016). After background correction, *M*_O2_ by fish was determined as μmol min^−1^ g^−1^ using standard equations for intermittent respirometry (Svendsen et al., 2016). All oxygen concentrations were corrected for salinity, barometric pressure, and temperature.

### SMR and RMR determination

We evaluated seven methods of estimating SMR (Chabot et al., 2016) using *M*_O2_ data collected between 20:00 and 06:00 in Trials 1 and 2, corresponding to 60 *M*_O2_ measurements per fish per trial: the mean of the lowest 10 data points (low10); the mean of the lowest 10% of the data, after removing the five lowest values (low10pc); quantiles that place SMR above the lowest 10-25% of the observations (q_0.1_, q_0.15_, q_0.2_, q_0.25_); and the mean of the lowest normal distribution (MLND). SMR estimated by low10 was lowest, although not statistically different from low10pc, q_0.1_, q_0.15_, or q_0.2_ (Table S1). In addition to providing a low value for SMR, the calculated value ought to agree with visual inspection of the raw data (Chabot et al., 2016). SMR values estimated by q_0.2_ and q_0.25_ best agreed with the distribution of *M*_O2_ from more trials than any other estimate. The analytical method also should be reproducible when applied to data generated from multiple trials with the same fish. SMR determined as low10pc, q_0.15_, and q_0.2_ were more highly correlated between Trial 1 and 2 (Pearson’s r > 0.80) than SMR determined by other methods (Pearson’s r < 0.80). As a final test of the robustness of SMR determination, we pooled all the data from 22 trials on 11 fish to generate a frequency distribution of 1320 *M*_O2_ values and then randomly sampled from this distribution to generate 1000 sets of 60 *M*_O2_ data points (as in the experimental trials). When SMR was calculated from these randomly generated datasets, q_0.2_ and q_0.25_ produced the fewest statistical outliers (Fig. S1). Only the q_0.2_ approach satisfied all of the criteria—it generated a low estimate of SMR; it agreed with the distribution of raw *M*_O2_ data; it was reproducible in repeated trials with the same fish; and it produced consistent values when applied to randomly generated datasets. Therefore, SMR determined by this approach was used for the remainder of these analyses. We also calculated routine metabolic rate (RMR), which includes spontaneous, uncontrolled activity in an otherwise quiet, post-absorptive fish, by taking the average of all 60 *M*_O2_ values collected between 20:00 and 06:00. Neither SMR nor RMR were determined for Trial 3 due to a limited number of *M*_O2_ measurements at normal air saturation. Hence, for *P*_crit_ determination in Trial 3 (see below), SMR or RMR for each fish was determined as its mean SMR or RMR from Trials 1 and 2.

### *P*_crit_ determination

We compared the following curve-fitting methods to describe *M*_O2_ as a function of *P*_O2_: broken-stick regression (BSR); nonlinear regression fit to a hyperbolic function, analogous to the Michalis Menten equation (MM); nonlinear regression fit to the Weibull function (W); and a linear function of *M*_O2_ measured at low *P*_O2_ (LLO). BSR was done using the Segmented package in R (Muggeo 2003). The nls() function of the base R package (R Core Team, 2017) was used to fit data to the MM and W functions. The MM function has the general form:

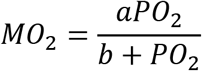

where *M*_O2_ is metabolic rate, *P*_O2_ is oxygen tension, and a and b are constants (*V*_max_ and *K*_M_, respectively, when applied to enzyme kinetics). The W function is:

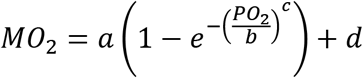

where *M*_O2_ is metabolic rate, *P*_O2_ is oxygen tension, and a, b, c, and d are constants. Because neither function has a parameter strictly equivalent to *P*_crit_, we report the value of b for the MM function (i.e., the *P*_O2_ when *M*_O2_ is 50% of the maximum *M*_O2_ extrapolated from that trial), and for both the MM and W functions, we determined the *P*_O2_ at which *M*_O2_ equals SMR or RMR. The last method (LLO) used the lm() function of the R base package (R Core Team, 2017) to fit a linear relationship between *M*_O2_ and *P*_O2_ to data collected after *M*_O2_ fell below that individual’s SMR. From this relationship, we determined the *P*_O2_ where *M*_O2_ equals SMR or RMR.

Importantly, SMR and RMR were determined during a previous overnight (~10 h) intermittent respirometry experiment, rather than from *M*_O2_ determined during the *P*_crit_ trial, when fish might become agitated and display increased *M*_O2_. In addition, BSR, M, and W used all the *M*_O2_ data collected during a given trial without subjective data elimination; LLO used only a subset of data (6-12 values) determined below the *P*_O2_ when *M*_O2_ fell below SMR. For all methods, the *P*_O2_ for a given *M*_O2_ was calculated as the mean *P*_O2_ over the measurement period (1 min for closed respirometry; 4 min for intermittent respirometry). Data for a representative fish, along with the methods for determining *P*_crit_, are shown in Figure 1.

**Fig. 1.**
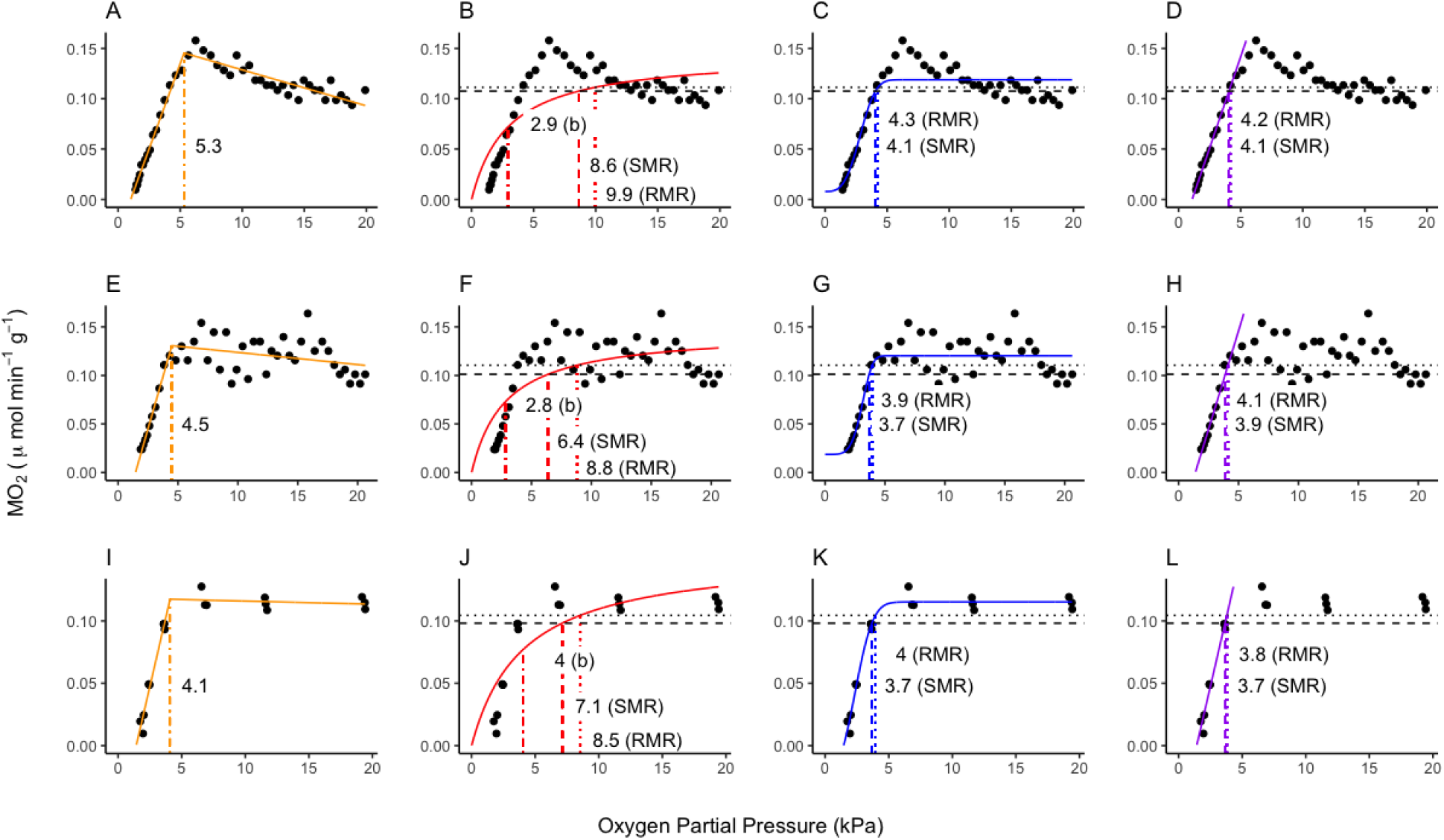
Model fits of each *P*_crit_ calculation method for a single *Fundulus grandis* used in three respirometry trials. Each row represents one experimental Trial: Fig. 1A-D, Trial 1 (closed respirometry); Fig. 1E-H, Trial 2 (closed respirometry); Fig. 1I-L, Trial 3 (intermittent respirometry). Each column represents one *P*_crit_ calculation methods: Fig. 1A, E, I, BSR where two linear segments were fit to the data (solid orange lines) and *P*_crit_ is the *P*_O2_ at their intersection (dashed orange line).; Fig. 1B,F,J, nonlinear regression using the M function (solid red line) and *P*_crit_ is the *P*_O2_ equal to *b* (analogous to *K*_M_ in enzyme kinetics), or *P*_O2_ when *M*_O2_ equals SMR or RMR; Fig. 1C,G,K, nonlinear regression using the W function (solid blue line) and *P*_crit_ is the *P*_O2_ when *M*_O2_ equals SMR or RMR; Fig. 1D,H,L, linear regression of *M*_O2_ versus *P*_O2_ at *M*_O2_ ≤ SMR (LLO method, solid purple line) and *P*_crit_ is the *P*_O2_ when *M*_O2_ equals SMR or RMR. For M, W, and LLO methods, SMR and RMR for this individual are shown by horizontal dashed and dotted lines, respectively. *P*_crit_ estimates are shown in the respective panels.

### Statistics

All statistical analyses were done in R v3.3.3 (R Core Team, 2017). The effects of analytical method (i.e., method used to calculate SMR or *P*_crit_) were determined within a given trial using linear mixed models (LMM) with analytical method as a fixed factor and fish as a random factor. All LMMs were fit using the lmer() function of the lme4 package (Bates et al., 2014) with *p*- values generated by the lmerTest package (Kuznetsova et al., 2017). All possible *post hoc* pairwise comparisons were made with *t*-tests on model fit means and employed *p*-values adjusted for false discovery using the emmeans package in R (Benjamini and Hochberg, 1995; Lenth 2018). Paired *t*-tests were used to compare of *P*_crit_ values based upon SMR and RMR within the MM, W, and LLO methods. The effects of respirometry method (closed versus intermittent) on the value of *P*_crit_ determined by a given analytical method were evaluated with LMM with respirometry method as a fixed factor and fish as a random factor. Correlation among values determined by a single analytical method across respirometry trials were evaluated with Pearson’s correlation coefficient (*r*).

## RESULTS AND DISCUSSION

### Models used to estimate *P*_crit_

The pattern of *M*_O2_ versus *P*_O2_ among fishes and other aquatic vertebrates has traditionally been modelled by the intersection of two straight lines (Yeager and Ultsch, 1989). In the present study, *P*_crit_ values estimated by BSR were among the highest and most variable estimates, including several that were >10 kPa (Fig. 2 and Table 1). In addition, *P*_crit_ values estimated by BSR were poorly reproducible between respirometry trials conducted with the same individuals under identical (closed respirometry) conditions (Table S2). These results are likely due to the variability of *M*_O2_ at levels of *P*_O2_ that do not limit oxygen uptake (i.e., at *P*_O2_ > *P*_crit_), as well as the tendency in some individuals for *M*_O2_ to increase as *P*_O2_ decreased from 20 to 5 kPa, resulting in a poor linear fit of *M*_O2_ data at high *P*_O2_ and influencing the intersection of two line segments. This variability occurred even though *P*_crit_ trials were conducted after > 24 h fasting, after 8-12 h since transferring fish to the respirometer, and during the dark phase of the photoperiod, when this species is less active. Owing to the variability of *M*_O2_ at high *P*_O2_, the use of BSR is frequently coupled with removal of *M*_O2_ data points that fail to meet certain criteria (see Claireaux and Chabot, 2016 and Wood, 2018 for examples). This practice has raised concern over the rationale and validity of applying data selection criteria (Claireaux and Chabot, 2016; Wood, 2018). In addition, direct comparisons of BSR with various non-linear regression approaches have shown that BSR is seldom the best model to fit *M*_O2_ data across a range of *P*_O2_ (Marshall et al., 2013; Cobbs and Alexander, 2018). Indeed, in a recent meta-analysis, BSR was the best model in only one out of 68 datasets fit with various statistical models (Cobbs and Alexander, 2018).

**Fig. 2.**
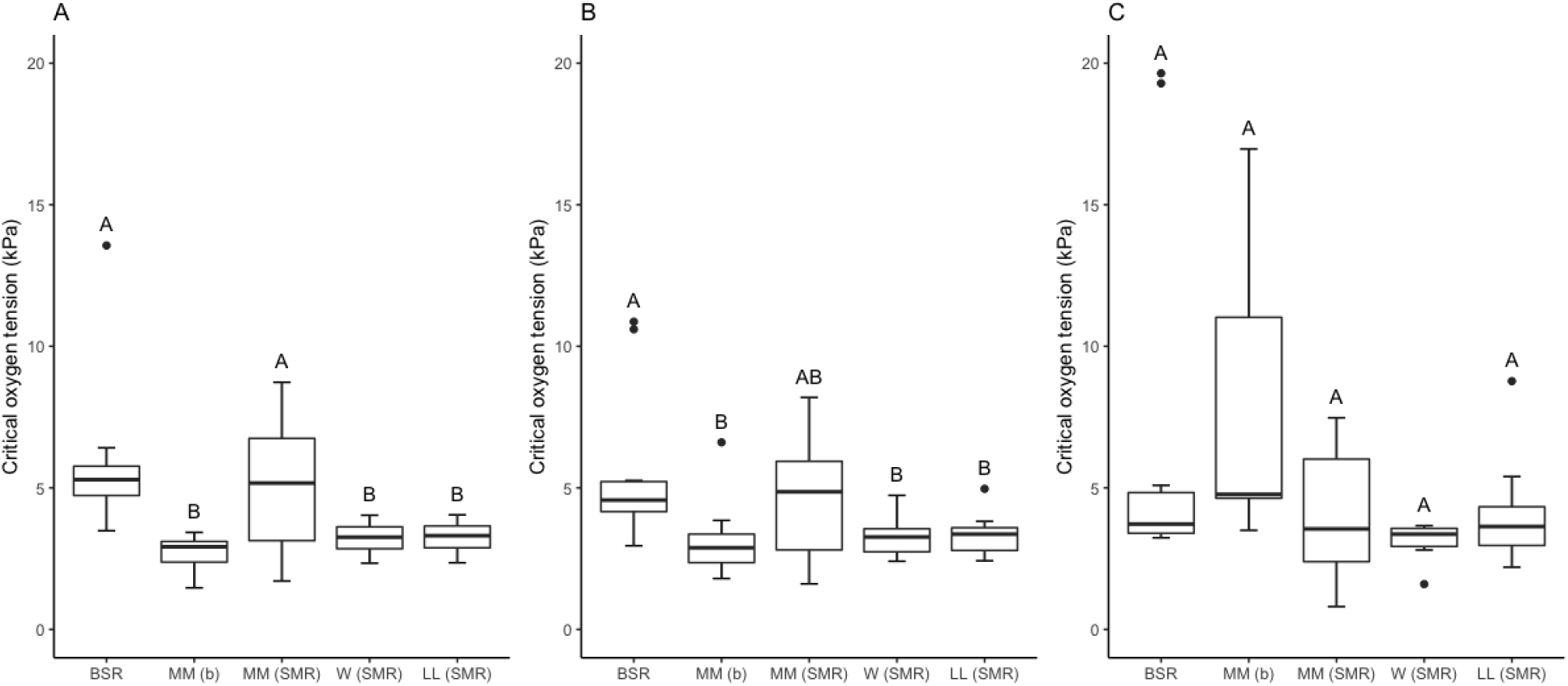
*P*_crit_ estimated by different analytical methods for *Fundulus grandis* in closed (Fig. 2A,B) and intermittent (2C) respirometry. Median values are indicated by the center line, upper and lower quartiles are upper and lower box boundaries, and the full data range are the whiskers (after removal of outliers, solid circles). *P*_crit_ estimates with different letters are significantly different within a trial (t-test, p < 0.05, false discovery corrected). Sample sizes (n) are 11 for each method in Fig. 2A, B, but varied among methods in Fig. 2C: BSR, n=11; MM(b), n=10; MM(SMR), n=9; W(SMR), n=6; LLO, n=11.

**Table 1.** Comparison of analytical method and respirometry format on the determination of *P*_crit_ (kPa; means ± SD) of the Gulf killifish, *Fundulus grandis*. Sample sizes (n) were 11, except where noted in parentheses.

|  | Trial 1 | Trial 2 | Trial 3 |
| --- | --- | --- | --- |
|  | (closed) | (closed) | (intermittent) |
| BSR | $5.8 \pm 2.7^a$ | $5.5 \pm 2.6^a$ | $6.6 \pm 6.4$ |
| MM (b) | $2.7 \pm 0.7^b$ | $3.1 \pm 1.3^b$ | $7.9 \pm 3.5^{**}(10)$ |
| MM (SMR) | $4.9 \pm 2.3^a$ | $4.5 \pm 2.1^{a,b}$ | $4.0 \pm 2.5(9)$ |
| W (SMR) | $3.2 \pm 0.5^b$ | $3.2 \pm 0.7^b$ | $3.1 \pm 0.8(6)$ |
| LLO (SMR) | $3.3 \pm 0.6^b$ | $3.3 \pm 0.7^b$ | $4.0 \pm 1.9$ |
| MM (RMR) | $8.5 \pm 4.6^*$ | $9.9 \pm 5.4^*$ | $5.2 \pm 3.1^*(8)$ |
| W (RMR) | $3.7 \pm 0.7^*$ | $3.8 \pm 0.6^*$ | $3.3 \pm 0.7^*(6)$ |
| LLO (RMR) | $3.6 \pm 0.7^*$ | $3.8 \pm 0.7^*$ | $4.7 \pm 2.3^*$ |

Means with different superscript letters are significantly different within a trial ( $t$ -test, $p < 0.05$ , false discovery corrected).
\*Significantly different from value estimated based upon SMR for the same analytical method within that trial (paired $t$ -test, $p < 0.05$ ).
\*\*Significantly higher than values determined by MM (b) method during closed respirometry (linear mixed model, $p < 0.05$ ).

With the advent and accessibility of nonlinear regression methods, it is possible to fit a variety of nonlinear functions to *M*_O2_ data. Here, we focussed on two nonlinear models, a hyperbolic function, analogous to the Michaelis-Menton equation for enzyme kinetics, and the Weibull function. Although the relationship between *M*_O2_ and *P*_*O2*_ in biological material as diverse as mitochondria to fishes can be hyperbolic (Tang, 1933; Gnaiger, 1993; Marshall et al., 2013), *M*_O2_ by *F. grandis* was poorly described by a hyperbolic function (Fig. 1). In addition, there is no consensus on what parameter of the MM function best describes the oxygen dependence of *M*_O2_. The parameter *b* is the *P*_O2_ when *M*_O2_ is half of the extrapolated maximum *M*_O2_ in that particular trial. Using *b* as an estimate of *P*_crit_ for closed respirometry yielded values that were reproducible among individuals (Fig. 2A,B), as well as between trials (Table S2), but were highly variable for intermittent respirometry (Fig. 2C). Also, it is not clear that this parameter has any particular meaning when applied to whole animal *M*_O2_, unlike its meaning in enzyme kinetics (Regan et al., 2019). In addition, the use of the parameter *b* assumes that the model fits the data well and that the upper asymptote of the MM function represents a definite, physiological maximum, neither of which were true in this study. Thus, we also used the equation of the hyperbolic function to estimate the *P*_O2_ when *M*_O2_ equals SMR, which resulted in high and variable estimates of *P*_crit_ (Fig. 2 and Table 1). Finally, the MM function also returned values which were either negative or above air-saturation in 10-20% of the datasets.

In their meta-analysis, Marshall et al. (2013) found that the Weibull function fit respirometric data better than other nonlinear functions, including the MM function. In the current study, the W function fit data from closed respirometry quite well, especially at low *P*_O2_ (Fig. 1). Like the MM function, though, there is no parameter of the W function that is analogous to *P*_crit_. Marshall et al. (2013) suggested that *P*_crit_ of a nonlinear function be estimated as the *P*_O2_ were the slope of the function approaches zero. In their analysis, the value of 0.065 was chosen as the slope giving a *P*_O2_ that “best approximates *P*_crit_”. This is a circular argument and requires prior knowledge of *P*_crit_, presumably based upon BSR. Rather than estimate an inflection point, we used the derived equation to determine the *P*_O2_ at which *M*_O2_ equalled SMR for each individual. For closed respirometry, this approach yielded highly reproducible values of *P*_crit_ similar to those determined by other methods in this study (Table S2). In contrast, for nearly half of the intermittent respirometry trials the W function failed to converge, severely limiting the usefulness of this approach. In addition, some software packages do not include nonlinear regression or they arrive at different solutions for the same dataset (personal observations).

For many fishes, the decline in *M*_O2_ at low *P*_O2_ is well described as a linear function of ambient oxygen, despite the variable and nonlinear relationship at higher *P*_O2_ (e.g., Claireaux and Chabot, 2016; Snyder et al., 2016). This was also the case in *F. grandis*, during both closed and intermittent respirometry (Fig. 1). Using the linear relationship between *M*_O2_ and *P*_O2_ at low *P*_O2_, *P*_crit_ was determined as the value of *P*_O2_ when *M*_O2_ equals SMR. This approach (LLO) yielded values similar to the MM method (based upon *b*), the W method, and previously published values for *F. grandis* (Virani and Rees, 2000). However, unlike the nonlinear methods, the LLO method successfully estimated *P*_crit_ values for all fish in all trials. In addition, this method is straight-forward and easy to implement, as long as SMR is accurately determined.

### An alternative to inflection point to determine P_crit_

With an equation for the relationship between *M*_O2_ on *P*_O2_, whether it be linear or nonlinear, it is possible to determine the value of *P*_O2_ for a specific value of *M*_O2_ rather than estimate an inflection point. In this approach, two critical issues must be addressed: the function must adequately describe the data and one must select the value of *M*_O2_ to interpolate. For many species, the relationship between *M*_O2_ and *P*_O2_ at low *P_O2_* values is well described by a straight line (current study; Affonso and Rantin 2005; Pan et al., 2016; Snyder et al., 2016; Thuy et al., 2010; Wong et al., 2017). With respect to the value of *M*_O2_ to use to solve for *P*_crit_, we and others advocate the use of SMR (Claireaux and Chabot, 2016). If oxygen drops below this level, the fish cannot sustain its minimal metabolic requirements via aerobic metabolism, thus representing a clear physiological limitation. Among fishes, RMR is more commonly used to determine *P*_crit_ (Rogers et al., 2016). This metabolic state includes routine, spontaneous activity, which may be more ecologically relevant than SMR (Fry and Hart, 1948; Rogers et al., 2016; Wood, 2018). For comparison, we also determined *P*_crit_ based upon RMR using the MM, W, and LLO functions (Fig. 1 and Table 1). Because RMR includes an undetermined level of activity, these estimates were significantly higher and generally more variable that *P*_crit_ based upon SMR. In addition, use of RMR complicates the interpretation of *P*_crit_ variation among individuals, experimental trials, or species: because this variation reflects differences in activity, comparisons of *P*_crit_ based upon RMR could obscure fundamental differences in oxygen extraction capacity. Indeed, Wong et al. (2017) found significant differences in *P*_crit_ among multiple species of Triggerfishes when using SMR to calculate *P*_crit_, but not when using RMR to estimate *P*_crit_.

### Recommendations

Based upon our results with *F. grandis* and the foregoing discussion, we propose that *P*_crit_ be defined as the *P*_O2_ where *M*_O2_ equals SMR. This recommendation requires that SMR be determined with high accuracy and using robust analytical techniques that yields a low value but is insensitive to occasional low outliers, agrees with the distribution of raw *M*_O2_ data, and is reproducible across multiple trials (Chabot et al., 2016). In the current experiments, the q0.2 method satisfied these criteria. Once SMR is determined, *P*_crit_ may then be determined in a continuation of the same experiment or in a different experiment if SMR is repeatable over time (Reemeyer et al., 2019). We recommend that *P*_crit_ be estimated as the *P*_O2_ where *M*_O2_ equals SMR based upon a linear relationship of *M*_O2_ and *P*_O2_ at low *P*_O2_ (i.e., the LLO method). The trial to determine *P*_crit_ can employ either closed or intermittent respirometry, as long as the experiment includes enough data points below SMR to provide a good linear fit. In the current study, *P*_crit_ deduced by the LLO method was lower, but not statistically so, when determined by closed respirometry compared to intermittent respirometry. Interestingly, although *P*_crit_ values were highly correlated between replicate trials of closed respirometry, they were not correlated between either trial of closed respirometry and the single trial of intermittent respirometry (Table S2). Both observations support the idea that respirometry method may influence *P*_crit_ (Regan and Richards, 2017; Snyder et al., 2016). Notwithstanding, the current study shows that method used to calculate *P*_crit_ is as important as respirometry format, highlighting the need to standardize analytical as well as experimental approaches in assessing the oxygen dependence of metabolism.

## Acknowledgements

We thank Mohammad Hamed for help with animal care.

## Data availability

Data and associated R script have been deposited at figshare.com (https://doi.org/10.6084/m9.figshare.8869253.v1).

## Supplementary Material

**Table S1:** Standard metabolic rate (μmol O_2_ min^−1^ g^−1^; means ± SD) of the Gulf killifish, *Fundulus grandis*, estimated by multiple calculation methods (see Materials and Methods and Chabot et al., 2016) from two respirometry trials using the same fish. For comparison, routine metabolic rate (RMR) was determined as the mean *M*_O2_ during each trial. Sample size = 11.

|  | Trial 1 | Trial 2 |
| --- | --- | --- |
| low10 | $0.092 \pm 0.012^a$ | $0.090 \pm 0.011^a$ |
| low10pc | $0.093 \pm 0.013^{a,b,c}$ | $0.092 \pm 0.012^{a,b}$ |
| q0.1 | $0.093 \pm 0.013^{a,b}$ | $0.091 \pm 0.011^{a,b}$ |
| q0.15 | $0.094 \pm 0.013^{a,b,c}$ | $0.093 \pm 0.012^{a,b,c}$ |
| q0.2 | $0.096 \pm 0.014^{a,b,c}$ | $0.096 \pm 0.012^{a,b,c}$ |
| q0.25 | $0.097 \pm 0.013^{b,c}$ | $0.096 \pm 0.012^{b,c}$ |
| MLND | $0.098 \pm 0.014^c$ | $0.099 \pm 0.014^c$ |
| RMR | $0.110 \pm 0.021^d$ | $0.116 \pm 0.021^d$ |

Means with different superscript letters are significantly different within a trial ( $t$ -tests on linear mixed model means, $p < 0.05$ false discovery corrected).

**Table S2:** Pearson’s correlation coefficients, *r*, comparing *P*_crit_ determined by various analytical techniques using data from multiple respirometry trials performed on the same individuals. Sample size are shown in parentheses (n).

|  | Trials 1 vs 2 | Trials 1 vs 3 | Trials 2 vs 3 |
| --- | --- | --- | --- |
| BSR | 0.18 (11) | -0.10 (11) | 0.34 (11) |
| MM (b) | 0.73* (11) | 0.54 (10) | 0.43 (10) |
| MM (SMR) | 0.83* (11) | -0.10 (9) | -0.35 (9) |
| W (SMR) | 0.75* (11) | 0.85* (6) | 0.82* (6) |
| LLO (SMR) | 0.74* (11) | -0.12 (11) | -0.29 (11) |
| MM (RMR) | 0.39 (11) | 0.57 (9) | 0.32 (8) |
| W (RMR) | 0.62* (11) | -0.09 (6) | -0.62 (6) |
| LLO (RMR) | 0.73* (11) | 0.14 (11) | -0.13 (11) |

\* $p < 0.05$

**Fig. S1.**
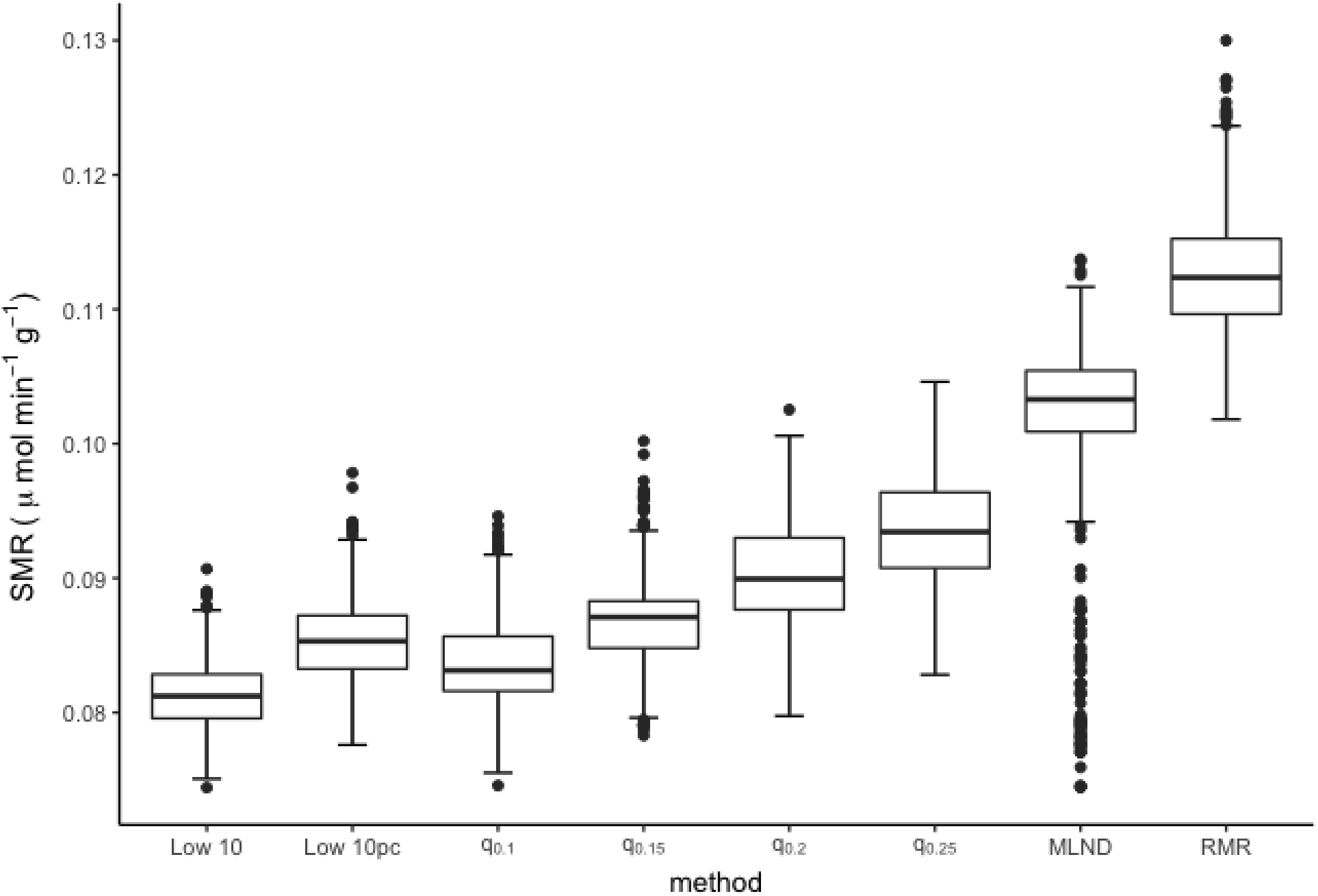
Standard metabolic rate (SMR) of *Fundulus grandis* calculated by different analytical methods from 1000 randomly generated datasets. All *M*_O2_ data from closed respirometry (1320 *M*_O2_ values measured across a range of *P*_O2_) were pooled and randomly sampled to generate 1000 sets of 60 *M*_O2_ values each. SMR was then calculated with the following methods: the mean of the lowest 10 data points (low10); the mean of the lowest 10% of the data after removing the 5 lowest points (low10pc); the 10 – 25% quantiles (q_0.1_, q_0.15_, q_0.2_, q_0.25_); and the mean of the lowest normal distribution (MLND) after fitting multiple normal distributions to the data (Chabot et al., 2016). For comparison, routine metabolic rate (RMR) was also calculated as the mean of all 60 *M*_O2_ values for each dataset. The whisker and box plots show the median (center line), upper and lower quartiles (upper and lower box boundaries), and full data range (whiskers) after removal of outliers (solid circles).

## REFERENCES

Affonso, E. G. and Rantin, F. T. (2005). Respiratory responses of the air-breathing fish Hoplosternum littorale to hypoxia and hydrogen sulfide. Comp. Biochem. Physiol. Part C Toxicol. Pharmacol. 141, 275–280.

Bates, D., Mächler, M., Bolker, B. and Walker, S. (2014). Fitting Linear Mixed-Effects Models using lme4. J. Stat. Softw. 67.

Benjamini, Y., Drai, D., Elmer, G., Kafkafi, N. and Golani, I. (2001). Controlling the false discovery rate in behavior genetics research. Behav. Brain Res. 125, 279–84.

Chabot, D., Steffensen, J. F. and Farrell, A. P. (2016). The determination of standard metabolic rate in fishes. J. Fish Biol. 88, 81–121.

Claireaux, G. and Chabot, D. (2016). Responses by fishes to environmental hypoxia: integration through Fry’s concept of aerobic metabolic scope. J. Fish Biol. 88, 232–251.

Cobbs, G. A. and Alexander, J. E. (2018). Assessment of oxygen consumption in response to progressive hypoxia. PLoS One 13, e0208836.

Diaz, R. J. and Rosenberg, R. (2008). Spreading dead zones and consequences for marine ecosystems. Science 321, 926–929.

Gnaiger, E. (1993). Adaptations to winter hypoxia in a shallow alpine lake. Ecophysiological energetics of Cyclops abyssorum and rainbow trout. Verh. Dtsch. Zool. Ges. 86, 43–65.

Hart, F. E. J. and Fry, T. S. (1948). The relation of temperature to oxygen consumption in the goldfish. Biol. Bull. 94, 66–77.

Kuznetsova, A., Brockhoff, P. B. and Christensen, R. H. B. (2017). lmerTest package: tests in linear mixed effects models. J. Stat. Softw. 82.

Lenth, R. (2019). emmeans: Estimated Marginal Means, aka Least-Squares Means. R package version 1.3.3. https://CRAN.R-project.org/package=emmeans

Marshall, D. J., Bode, M. and White, C. R. (2013). Estimating physiological tolerances - a comparison of traditional approaches to nonlinear regression techniques. J. Exp. Biol. 216, 2176–2182.

Muggeo, V. M. R. (2003). Estimating regression models with unknown break-points. Stat. Med. 22, 3055–3071.

Pan, Y. K., Ern, R. and Esbaugh, A. J. (2016). Hypoxia tolerance decreases with body size in red drum Sciaenops ocellatus. J. Fish Biol. 89, 1488–1493.

Rabalais, N. N., Díaz, R. J., Levin, L. A., Turner, R. E., Gilbert, D. and Zhang, J. (2010). Dynamics and distribution of natural and human-caused hypoxia. Biogeosciences 7, 585–619.

Reemeyer, J. E., Harris, J. C., Hernandez, A. M. and Rees, B. B. (2019). Effects of passive integrated transponder tagging on cortisol release, aerobic metabolism and growth of the Gulf killifish Fundulus grandis. J. Fish Biol. 94, 422–433.

Regan, M. D., Mandic, M., Dhillon, R. S., Lau, G. Y., Farrell, A. P., Schulte, P. M., Seibel, B. A., Speers-Roesch, B., Ultsch, G. R. and Richards, J. G. (2019). Don’t throw the fish out with the respirometry water. J. Exp. Biol. 222, jeb200253.

Regan, M. D. and Richards, J. G. (2017). Rates of hypoxia induction alter mechanisms of O 2 uptake and the critical O 2 tension of goldfish. J. Exp. Biol. 220, 2536–2544.

Rogers, N. J., Urbina, M. A., Reardon, E. E., McKenzie, D. J. and Wilson, R. W. (2016). A new analysis of hypoxia tolerance in fishes using a database of critical oxygen level (P_crit_). Conserv. Physiol. 4, cow012.

Rosewarne, P. J., Wilson, J. M. and Svendsen, J. C. (2016). Measuring maximum and standard metabolic rates using intermittent-flow respirometry: a student laboratory investigation of aerobic metabolic scope and environmental hypoxia in aquatic breathers. J. Fish Biol. 88, 265–283.

Snyder, S., Nadler, L. E., Bayley, J. S., Svendsen, M. B. S., Johansen, J. L., Domenici, P. and Steffensen, J. F. (2016). Effect of closed v. intermittent-flow respirometry on hypoxia tolerance in the shiner perch Cymatogaster aggregata. J. Fish Biol. 88, 252–264.

Svendsen, M. B. S., Bushnell, P. G., Christensen, E. A. F. and Steffensen, J. F. (2016). Sources of variation in oxygen consumption of aquatic animals demonstrated by simulated constant oxygen consumption and respirometers of different sizes. J. Fish Biol. 88, 51–64.

Tang, P.-S. (1933). On the rate of oxygen consumption by tissues and lower organisms as a function of oxygen tension. Q. Rev. Biol. 8, 260–274.

Thuy, N. H., Tien, L. A., Tuyet, P. N., Huong, D. T. T., Cong, N. V., Bayley, M., Wang, T. and Lefevre, S. (2010). Critical oxygen tension increases during digestion in the perch Perca fluviatilis. J. Fish Biol. 76, 1025–1031.

Ultsch, G.R., Jackson, D.C. and Moalli, R.J. (1981) Metabolic oxygen conformity among lower vertebrates—the toadfish revisited. J. Comp. Physiol. 142, 439–443.

Virani, N. A. and Rees, B. B. (2000). Oxygen consumption, blood lactate and inter-individual variation in the gulf killifish, Fundulus grandis, during hypoxia and recovery. Comp. Biochem. Physiol. - A Mol. Integr. Physiol. 126, 397–405.

Wong, C. C., Drazen, J. C., Callan, C. K. and Korsmeyer, K. E. (2018). Hypoxia tolerance in coral-reef triggerfishes (Balistidae). Coral Reefs 37, 215–225.

Wood, C. M. (2018). The fallacy of the Pcrit – are there more useful alternatives? J. Exp. Biol.221, jeb163717.

Yeager, D. P. and Ultsch, G. R. (1989). Physiological regulation and conformation: a BASIC program for the determination of critical points. Physiol. Zool. 62, 888–907.

